# Scalable inference of transcriptional kinetic parameters from MS2 time series data

**DOI:** 10.1101/2020.12.04.412049

**Authors:** Jonathan R. Bowles, Caroline Hoppe, Hilary L. Ashe, Magnus Rattray

## Abstract

**Motivation:** The MS2-McP (MS2 coat protein) live imaging system allows for visualisation of transcription dynamics through the introduction of hairpin stem-loop sequences into a gene. A fluorescent signal at the site of nascent transcription in the nucleus quantifies mRNA production. computational modelling can be used to infer the promoter states along with the kinetic parameters governing transcription, such as promoter switching frequency and polymerase loading rate. However, modelling of the fluorescent trace presents a challenge due its persistence; the observed fluorescence at a given time point depends on both current and previous promoter states. A memory-adjusted Hidden Markov Model (mHMM) was recently introduced to allow inference of promoter activity from MS2-McP data. However, the computational time for inference scales exponentially with gene length and the mHMM is therefore not currently practical for application to many eukaryotic genes.

**Results:** We present a scalable implementation of the mHMM for fast inference of promoter activity and transcriptional kinetic parameters. This new method can model genes of arbitrary length through the use of a time-adaptive truncated compound state space. The truncated state space provides a good approximation to the full state space by retaining the most likely set of states at each time during the forward pass of the algorithm. Testing on MS2-MCP fluorescent data collected from early *Drosophila melanogaster* embryos indicates that the method provides accurate inference of kinetic parameters within a computationally feasible timeframe. The inferred promoter traces generated by the model can also be used to infer single-cell transcriptional parameters.

**Availability:** Python implementation available at https://github.com/ManchesterBioinference/burstInfer, along with code to reproduce the examples presented here.

## 1 Introduction

Recent advances in *in vivo* live imaging technologies (Pichon *et al.*, 2018) have created a pressing need for algorithms capable of analysing large, complex biological datasets. Live imaging techniques, such as the MS2-MCP system, have been of particular interest to the developmental biology community due to the ability to visualise transcription at singlecell resolution *in vivo*. As correct spatial and temporal control of gene expression is of fundamental importance during both normal development and disease, the ability to analyse the rich datasets generated by live imaging approaches is vital.

The MS2-MCP system allows for the quantification of transcription in real-time through the introduction of hairpin structures into a gene of interest (Pichon *et al.*, 2018). Following the entry of the promoter into an active state, elongation of RNA Polymerase II (Pol II) along the gene body results in the production of nascent mRNA transcripts containing hairpin stem-loops. Binding of the MCP fluorescent protein to this hairpin structure allows for detection of the resulting fluorescent signal by fluorescence microscopy (Bertrand *et al.*, 1998; Garcia *et al.*, 2013; Lucas *et al.*, 2013). Quantification of this fluorescent signal results in a fluorescent time series, which acts as a proxy for transcriptional output at each transcription site (Lucas *et al.*, 2013; Garcia *et al.*, 2013; Bertrand *et al.*, 1998). The ability to track the fluorescence of accumulated nascent mRNA at transcription foci (and therefore levels of transcriptional activity) over time and at single-cell resolution opens up the possibility of investigating spatial and temporal transcriptional dynamics in model organisms, in addition to the response of tissue culture cells to external stimuli (Pichon *et al.*, 2018). The use of the MS2-MCP system allows for the collection of temporal transcriptional data, an advantage over the static ‘snapshots’ of transcription generated using techniques such as single molecule fluorescent in *situ* hybridisation (smFISH) (Pichon *et al.*, 2018).

Transcription is now understood to be a highly dynamic process, with many genes producing transcripts in discrete pulses, or ‘bursts’, of transcriptional activity (Coulon *et al.*, 2013; Raj *et al.*, 2008; Chubb *et al.*, 2006; Golding *et al.*, 2005);. Transcriptional bursting has been observed in organisms ranging from Drosophila to vertebrates and is implicated in both normal development and disease (Raj *et al.*, 2008; Eldar *et al.*, 2010); bursting is of particular interest to the gene regulation community, as many key developmental genes appear to exhibit bursting-like behaviour (Lenstra *et al.*, 2016). Mathematical modelling of transcriptional bursting may be described by a set of kinetic parameters which report the frequency, amplitude and duration of transcriptional bursts (Zoller *et al.*, 2018; Li *et al.*, 2018; Dar *et al.*, 2012; Raj *et al.*, 2006; Fukaya *et al.*, 2016; Corrigan *et al.*, 2016). Previous work on mathematical modelling of transcriptional bursting has focused on inference of these transcriptional parameters through analysis of either static smFISH snapshots (Mueller *et al.*, 2013; Bahar Halpern *et al.*, 2015; So *et al.*, 2011; Gómez-Schiavon *et al.*, 2017) or MS2-MCP time series data (Corrigan *et al.*, 2016; Garcia *et al.*, 2013; Fukaya *et al.*, 2016; Berrocal *et al.*, 2018; Lammers *et al.*, 2020; Tantale *et al.*, 2016; Bothma *et al.*, 2014). The ability to infer these kinetic parameters opens up the possibility of providing a deeper insight into the spatio-temporal regulation of bursting at single-cell resolution.

While MS2-MCP time series data allows for visualisation of nascent transcription at single-cell resolution in real-time, inference of kinetic parameters from MS2-MCP data presents a number of unique challenges (Gregor *et al.*, 2014). Crucially, the presence of persistent fluorescence within the signal complicates inference of transcriptional kinetic parameters (Corrigan *et al.*, 2016; Lammers *et al.*, 2020). Upon the promoter entering an active state, RNA Polymerase (Pol II) commences elongation along the gene body, leading to a fluorescent signal through MCP-fluorescent protein binding. When the promoter becomes inactive, the fluorescent signal does not immediately cease. Pol II molecules are still in transit along the gene body and the incomplete mRNA transcripts are bound by MCP-fluorescent proteins. Inference of kinetic parameters therefore requires an algorithm capable of taking this persistence into account.

Previous work (Lammers *et al.* (2020)) incorporated the persistence of the MS2 signal through implementing a memory-adjusted hidden Markov model (mHMM), building on an earlier hidden Markov model for MS2-GCP parameter inference (Corrigan *et al.* (2016)). The transition probabilities and emission values of the model correspond to the promoter switching frequencies and Pol II loading rate, respectively, which together are sufficient to describe the bursting dynamics of the system. From an initial active or inactive state (π), the promoter switches between active and inactive states according to the transition matrix, loading polymerase onto the gene while in the active state at a rate determined by the model emission parameter (Figure 1A).

**Figure 1:**
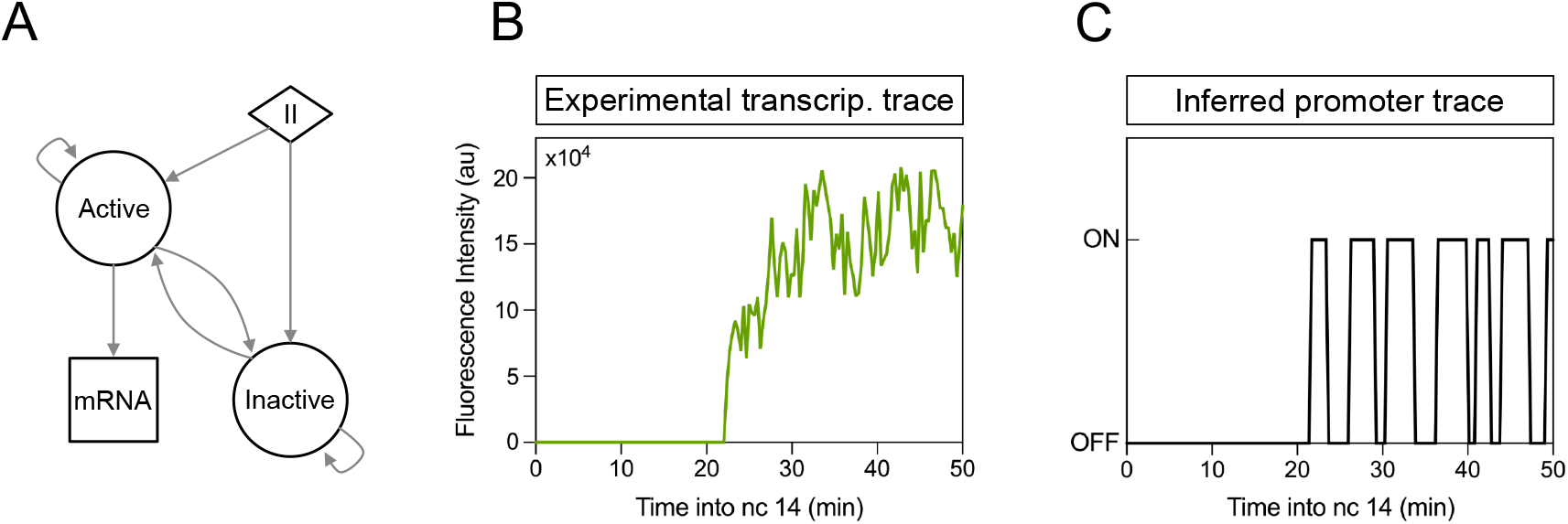
The model structure and basic principle behind *burstInfer*. **A**: Dynamic memory-adjusted hidden Markov model state diagram. At the beginning of the time sequence the promoter is in either the active or inactive state (π). Over the course of the time series the promoter switches stochastically between the active and inactive states according to the *k_on_* and *k_off_* burst parameters. While in the active state Pol II molecules are loaded onto the gene and mRNA transcripts are produced at a rate determined by the model emission parameter. **B**: Example MS2 fluorescence time series trace for a single nucleus in a *Drosophila* embryo showing nascent *ush* transcription. **C**: The promoter sequence inferred by the model corresponding to the fluorescent trace in B. These promoter traces can be used to generate single-cell parameters.

Persistence in the signal is dealt with through the inclusion of a window parameter, *W*, that models the dependence of the recorded fluorescence on the previous W promoter states, each of which may take one of *K* (here 2) values. The inclusion of the window parameter results in *K^W^* compound states to fully describe the system. This exponential scaling becomes problematic when dealing with long genes, as the dependence of the window parameter on elongation time (and therefore gene length) may lead to infeasible computational times.

In this paper we present a modified form of the mHMM, the Dynamic Memory-Adjusted Hidden Markov Model (dmHMM), referred to as *burstInfer*, for fast inference of kinetic parameters from MS2-MCP data. The algorithm represents a significant speed boost over the original mHMM technique when applied to long genes, removing the exponential time-scaling of the technique with gene length. Results indicate that the use of a reduced compound state space is sufficient to accurately infer kinetic parameters relative to the original model, while significantly reducing computational time for longer genes, making inference of kinetic parameters for genes of all sizes feasible.

## 2 Methods

### 2.1 Model Formulation

Following insertion of the MS2 stem-loop sequences into the gene of interest, elongation of Pol II along the length of the gene body results in the generation of a fluorescent time series signal. We intend to model the dynamics of these recorded fluorescent signals, with the aim of extracting the kinetic parameters driving expression of the target gene. Following the mHMM formulation derived by Lammers *et al.*, whose model this paper extends, we denote an individual fluorescent signal (corresponding to one transcription foci) as *y* = {*y_1_, y_2_…,y_T_*}, with *T* denoting the number of time points within the individual trace (Figure 2). We assume that the promoter may be in one of *K* = 2 effective states, i.e. active or inactive. The promoter switches between hidden states *z* at time step *t* according to the *K* × *K* transition matrix, *A* = *p*(*z_t_* |*z_t-1_*). *A_kl_* represents the probability of making the transition from hidden promoter state *k* to hidden promoter state *l* during time step *t*. Transitions between hidden promoter states *z_t_* are assumed to satisfy the Markov property, i.e. the hidden promoter state at a given time point depends only upon the hidden promoter state at the previous time point (Lammers *et al.*, 2020).

**Figure 2:**
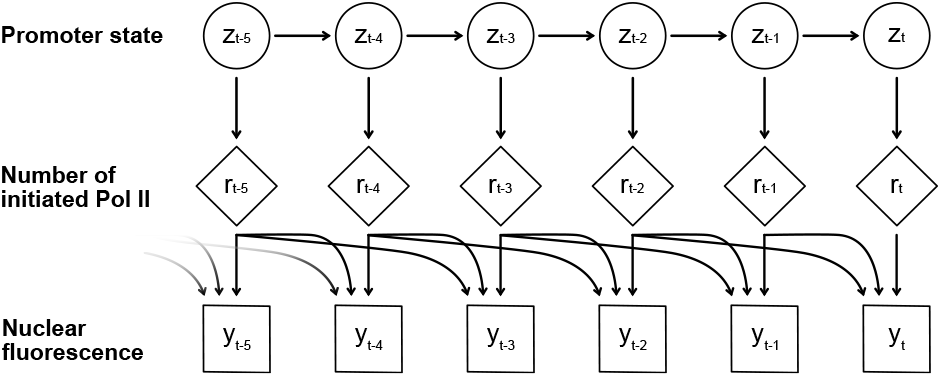
Diagram illustrating the dependence of the measured fluorescent signal at the present time, *t*, on both the present promoter state and previous promoter states falling within the observation time window, *W*. This timedependence arises due to the persistence in the MS2 signal caused by Pol II still being in transit down the gene body following the promoter becoming inactive. The example shown here is for window size W=3.

Each effective state *z_t_* is associated with a polymerase initiation rate, *r(k)*, representing the number of Pol II molecules loaded onto the gene in a given minute. The fluorescence data presented here are shown in terms of arbitrary units of fluorescence. Quantification of the transcriptional output of cells using smFISH may be used to calibrate the signal in terms of Pol II number instead (Garcia *et al.*, 2013; Lammers *et al.*, 2020; Hoppe *et al.*, 2020). The maximum fluorescence emission per time step *t* for each effective state is defined as *v(k)* = *Fr(k)*, where *F* is a calibration factor used to convert the units of arbitrary fluorescence to units of Pol II (Lammers *et al.*, 2020).

The recorded fluorescence intensity at a given time point (Figure 1B) depends upon not only the fluorescence generated during the previous time step, but also the cumulative fluorescence generated by Pol II in transit along the length of the gene during previous time steps. To model this dependence upon previous time steps the concept of a sliding window, *W*, is introduced into the model. This window, or memory, represents the dependence of the observation *y_t_* at time point t on not only the hidden promoter state *z_t_* at the current time point but also the previous *W* hidden promoter states (depicted in Figure 2). The value of *W* is gene-dependent and is calculated as 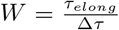, where *τ_elong_* is the elongation time and Δτ is the size of an individual time step, i.e. the time resolution of the data. Hidden promoter states falling outside the previous *W* time points can be assumed not to contribute to the recorded fluorescence at time point *t*, as Pol II initiated at that particular time point is no longer in transit along the gene.

To model this dependency of the observed fluorescence at time point t on the previous *W* hidden promoter states *z_t_*, the concept of a compound state *s_t_* = {*z_t_, z_t-1_*,…, *z_t-W+1_*} is introduced. *s_t_*, a 1 × *W* vector, encodes the sequence of *W* hidden promoter states up to and including the current hidden promoter state at time point *t*. At each given time point the previous *W* — 1 promoter states are deterministically passed to the new compound state, becoming the 1…*W* - 1 elements of the new compound state vector, with the *W^th^* compound state at time point *t* being determined stochastically by the state transition matrix *A*. In the original mHMM model, each compound state takes one of *K^W^* different values, as each of *W* hidden promoter states may take one of *K* values (Lammers *et al.*, 2020). This exponential scaling with window size *W* imposes a significant computational burden. How our model addresses this is detailed in the following section. As in the original mHMM model, the emissions of the Hidden Markov Model are described by a Gaussian distribution with mean *μ* and standard deviation *σ*. The initial hidden promoter states at time *t* = 0 are given by a 1 × *K* vector *π*. The joint distribution of compound states and observed fluorescence values is given by:

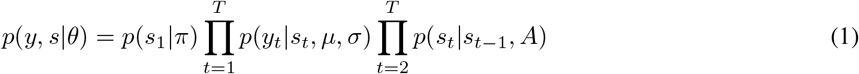

Expectation Maximisation is used to infer the Hidden Markov Model parameters, 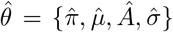. The use of an approximate inference technique renders inference of the model parameters computationally tractable. However, the exponential scaling of computation time with window size represents a significant problem for longer genes.

### 2.2 Dynamic State Space Truncation

In order to circumvent the exponential scaling of the algorithm with window size we propose a dynamic reduced state space variant of the mHMM, which uses a truncated state space to avoid exponential scaling in computational time. We illustrate the advantages of the dmHMM using a specific example implementation of the mHMM model with *K* = 2 promoter states and a window size of 19, as would be required to model the Drosophila melanogaster gene *u*-shaped (ush), which is 16825 base pairs in length (isoform C). Nascent transcription was captured at 20s time resolution. This results in a compound state which may take on K^W^ = 2^19^ = 524288 values. Repeated manipulation of the resulting K^W^ × *t* state matrix while performing expectation maximization requires a significant amount of computational time, which cannot be improved significantly by increasing available computational power.

The required computational time may be reduced by observing that although 524288 possible compound state values are required to fully specify the model, the majority of these compound states will have very low (often negligible) associated probability values, and can therefore be excluded from the model without impacting predictive performance. For example, during portions of the fluorescence signal recorded during the initiation of a transcriptional burst, compound states associated with inactive promoter states during the initial part of the compound state and active promoter states during the latter part of the compound state would be much more likely than compound states with sequences of promoter states associated with a very different observed fluorescence pattern, e.g. falling fluorescence levels or sustained inactivity.

Truncation in the model is enforced through the use of an allowed memory, *M*, with *M* < *K^w^*. *M* is selected so as to reduce computational time without significantly impacting the performance of the algorithm. The use of *M* results in a reduced promoter state space, Φ_t_, replacing s and reducing the scaling of the forward algorithm with window size from exponential to linear scaling. To select a set of *M* likely compound states at time *t* + 1 the forward algorithm is used to rank the 2M next possible states starting from *M* at time *t*. The forward algorithm computes the probability of the data up to the current time and being in each state, therefore the most likely states can be prioritized and the least likely are removed from the model until *M* distinct compound states remain.

An example of model truncation using a single trace of *ush* MS2 data is shown in Figure 3, with an allowed memory of 4 states specified for illustration purposes. Each box represents an individual state, with the leftmost number giving the binary representation of the promoter state (1 for on and 0 for off) and the rightmost number giving the log forward variable associated with each state. The state space expands during the forward algorithm until the allowed value of *M* is reached at *t* = 1. Forward variables are calculated for each allowed transition (the previous promoter state with either a 0 or 1 inserted at the rightmost bit) and are ranked. The least likely forward variables are eliminated (red outline), with the most likely states becoming the new reduced state space (blue outline). The process is repeated until the end of the trace (here t=3).

**Figure 3:**
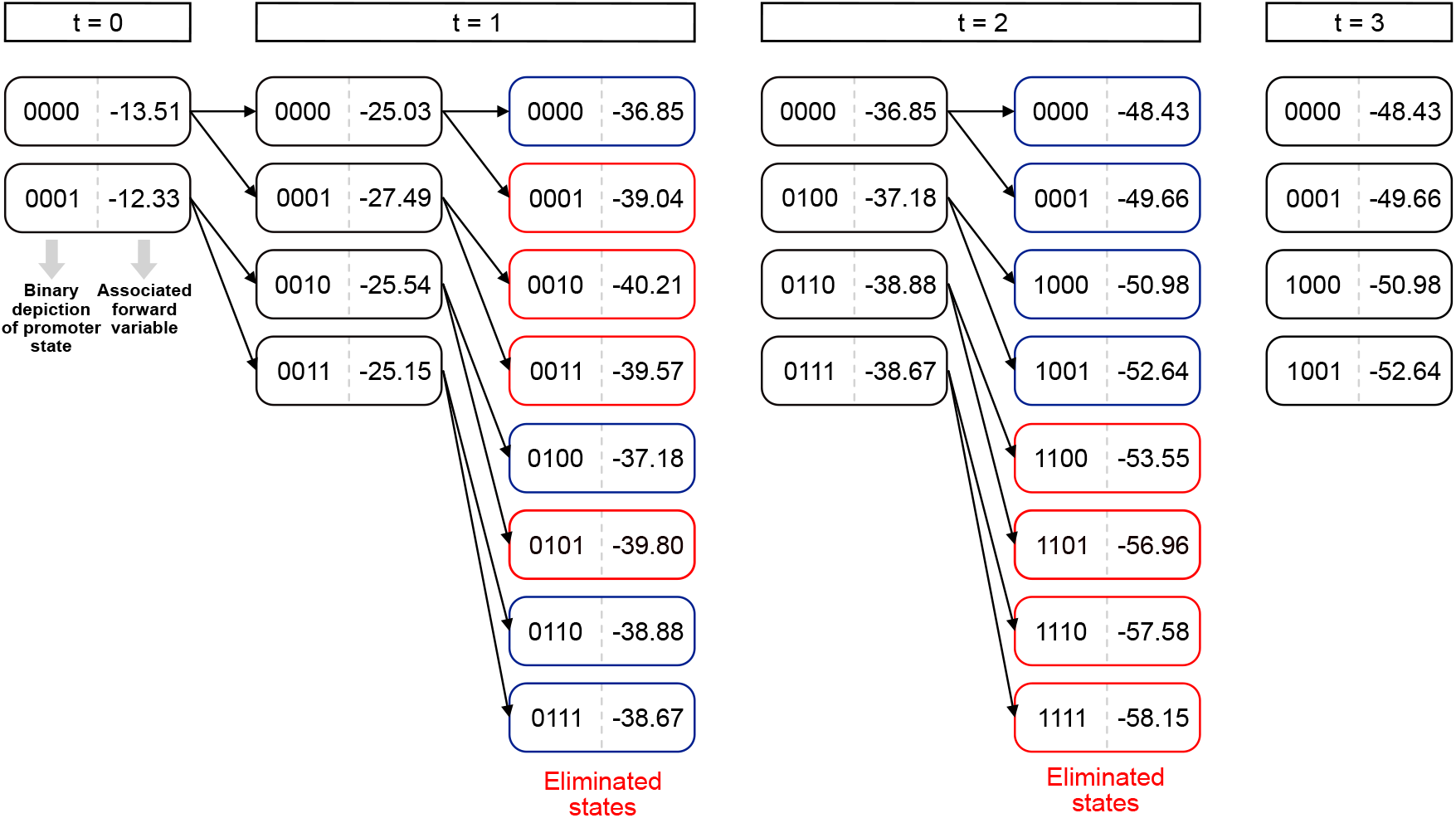
Example illustrating state-space truncation carried out as part of the HMM forward algorithm, using example data derived from the *Drosophila ush* gene. Each oblong bubble represents a compound promoter state at a particular time point with the number on the left representing the binary representation of the promoter state and the number on the right showing the log probability associated with each forward variable. The promoter starts at time *t* = 0 in either the inactive (0000) or active (0001) state (the rightmost bit indicates the current state). At time *t* = 1 the promoter can switch to either of two states from each of these two states, causing the state space to expand from 2 to 4 possible compound states (i.e. inactive to inactive, inactive to active, active to inactive, active to active). At time *t* = 2 the possible state space doubles again to 8 compound states. At this point truncation is carried out - the compound states are ranked according to probability and the least likely states are eliminated. The number of eliminated/retained states is determined by the window size - a window value of W = 2, giving a number of allowed states of 2^2^ is shown here so that elimination can be visualised. In practice the highest number of allowed states that is computationally feasible is used instead. This process of truncation and elimination is carried out until the end of each trace contained in the entire dataset. This truncated graph then becomes the state space for the entire model.

### 2.3 Inferring Single-cell Transcriptional Parameters

In addition to inferring ‘global’ model estimates for burst amplitude, frequency and duration for a given dataset, our model can be used to infer single-cell transcriptional parameters, i.e. burst parameters for each individual cell within the expression domain, rather than a global estimate for the entire expression domain or region of interest.

Training the model using the forward-backward algorithm yields estimates of 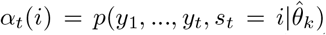, the joint estimate of the observed fluorescence up to time t and the compound hidden promoter state at time *t* and 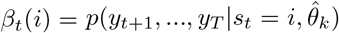, the conditional probability of the observations from (*t* + 1) to the end of each trace, given the current hidden promoter state. Combining these variables with the expression for the likelihood of the observed fluorescence values given the model parameters, 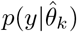, gives the following:

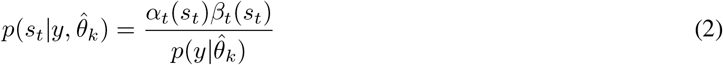

where 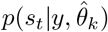 denotes the probability of the promoter being in an active or inactive state at a given time point *t*, given the observed fluorescence and inferred model parameters. Taking the argmax of Equation (2) at each time point gives a sequence of the most likely promoter states at each observed time step. As previously mentioned, the *Drosophila* gene *ush* is used here as an example. MS2 stem-loops were inserted into the endogenous *ush* gene 5’UTR region, allowing us to visualise transcription in the form of nascent MCP-GFP fluorescence (Figure 1B). The inferred promoter trace calculated using Equation (2) corresponding to this time series is shown in Figure 1C.

In addition to providing a way of visualising the model fit, these inferred promoter traces may be used to calculate single-cell transcriptional parameters, so that in addition to giving single maximum likelihood parameters estimates for a given dataset, i.e. a *k_on_*, *k_off_* and emission term for the set of traces used to train the model, each cell in the expression domain is assigned each of these parameters. An example of these parameters using *Drosophila u-shaped* data from Hoppe *et al.*, 2020 is shown in Figure 5 (details of the dataset used are given in Section 3.3).

**Figure 5:**
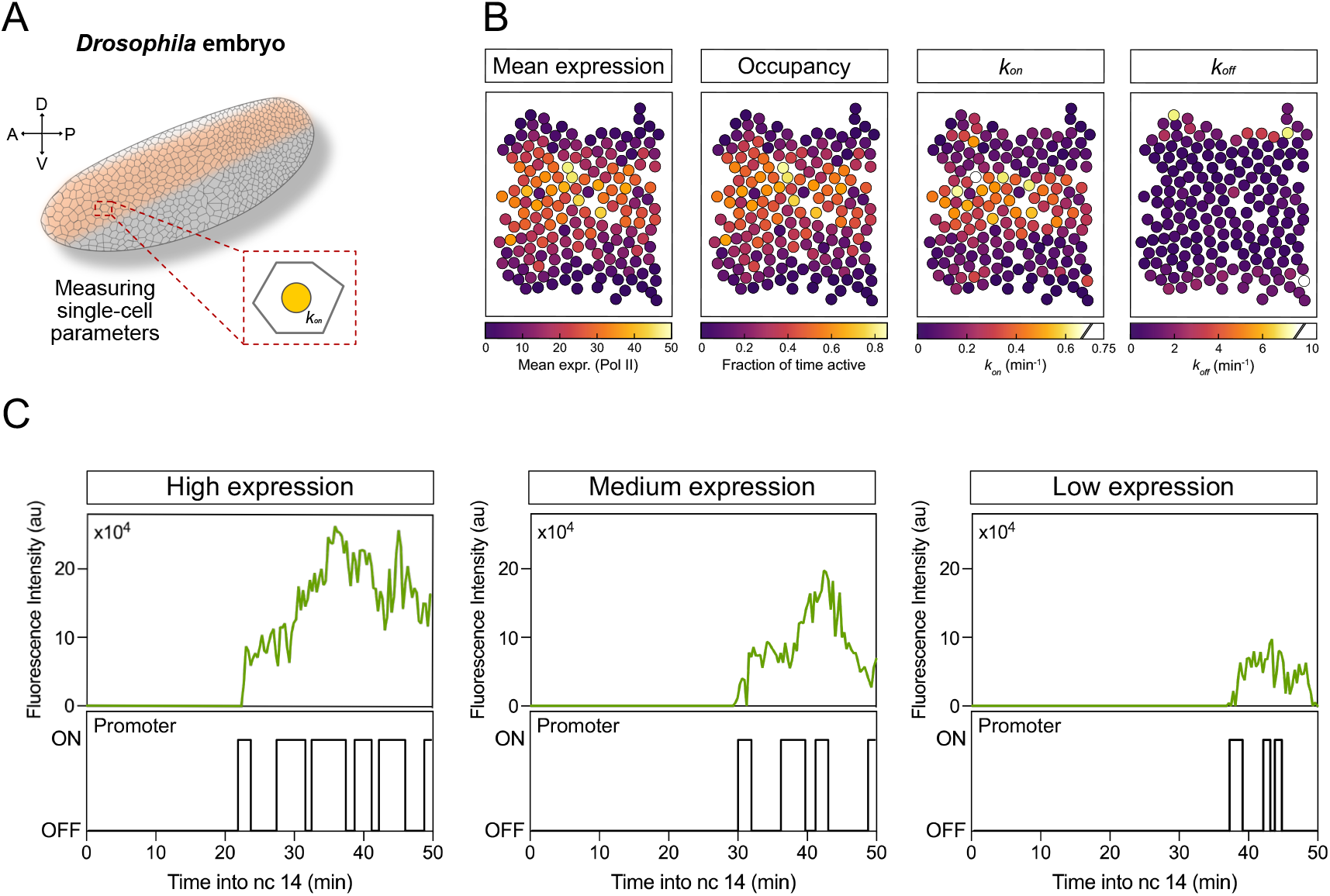
Example inferred single-cell parameter using *Drosophila ush* data from Hoppe *et al.* (2020). **A**: The expression domain of the *ush* gene shown in the cartoon was divided into three separate regions, corresponding to high, medium and low levels of expression, with the model trained separately on each of these three regions. The inferred global parameters for each region were used to infer the most likely promoter path corresponding to each fluorescent trace. **B**: Heatmaps of the measured mean expression level, along with the *k_on_, k_off_* and Occupancy 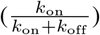 parameters for each cell are shown. Analysis of single-cell parameters in this case revealed *k_on_* as the main determinant of expression level. **C**: Example fluorescent traces and corresponding inferred promoter paths for each of the three regions. See Hoppe *et al.* (2020) for further details.

The calculation of the transition parameters is achieved through a simple counting-based technique, where the number of normalised on-to-off and off-to-on transitions is counted from the inferred promoter traces. These counts are used to create transition matrices for each trace, which are then converted to transition rates (in a similar way to the calculation of the global parameters). The single-cell emission term is a reduced form of the emission term from the global model (see Lammers *et al.*, 2020):

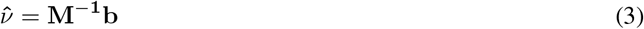

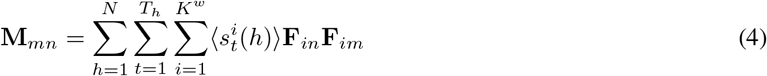

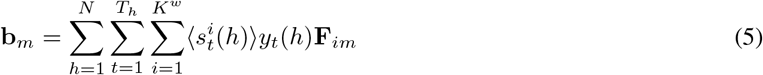

where the 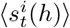 term becomes a delta function due to the state probabilities already being known.

## 3 Results

### 3.1 Comparison of Inferred Parameters

To demonstrate the ability of the truncated model to approximate the results obtained using the full hidden Markov model, we created synthetic fluorescent traces for a gene of window size 11 and tested convergence between the truncated and full models for this dataset (Figure 4A). The relative error between the full and truncated models falls smoothly as the state space of allowed states is increased, with the relative error falling to less than 1% at *M* = 128 where the size of the full model here would be 2^11^ = 2048 compound states.

**Figure 4:**
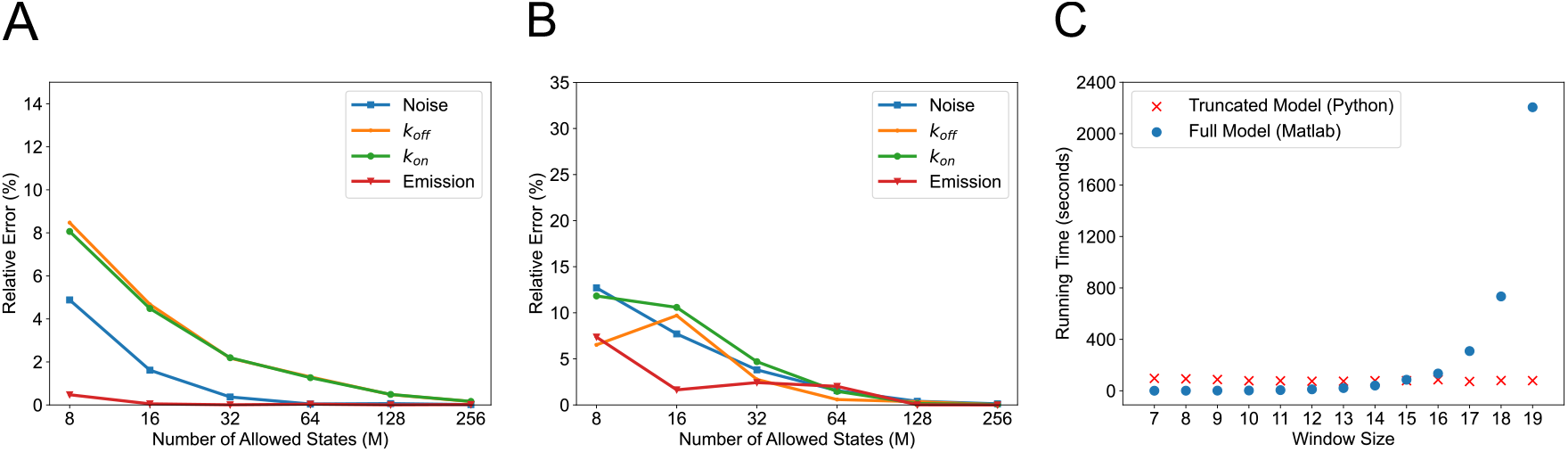
Verifying the model fit to real and synthetic data and visualising the non-exponential relationship between window size and running time. **A**: Plot of the relative error between the truncated and full model for 4 parameters as a function of an increasing number of allowed states. As the number of allowed states, *M* is increased the truncated model converges towards the full model. A Markov Process was used to generate synthetic MS2 data corresponding to a window size of 11. 2^11^ = 2048 compound states would be necessary to fully describe the data under the full model. By using a reduced subset of only 128 states, the relative error between the full and truncated models is reduced to less than 1%. Subfigure **B**: plot of the relative error between the truncated and full models for real *Drosophila hnt* data. The number of compound states necessary to fully specify the *hnt* model is 512. *hnt* has been chosen as inference using the full model is still possible for this relatively short gene. Subfigure **C**: running time for a single EM step in seconds for both models. For shorter window sizes the original model is faster due to decreased overhead from not having to calculate least likely states etc. For longer window sizes the exponential scaling of the original algorithm becomes an issue.

To test the model on experimental MS2-MCP data where is it possible to fit the full model, both the full and truncated models were trained on a dataset of MS2 fluorescent traces for the *Drosophila melanogaster gene hindsight (hnt)*. The hnt gene has length of 7441 base pairs between the MS2 probe and the end of the gene body, in conjunction with an MS2 cassette length of 1290 base pairs, a window size of 9 was specified. The results of training the model using both the full and truncated models can be seen in Figure 4B, a plot of relative error between the truncated and full (‘true’) model parameters as a function of increasing number of allowed states. Each curve represents a separate parameter of the model. The model was trained specifying 50 separate runs of expectation-maximisation for each value of *M*. The convergence of the truncated model parameters to the full model parameters is apparent from the diagram.

### 3.2 Comparison of Computational Time

Next, we compared the scaling of computational time for a single step of the expectation maximisation algorithm for the truncated model and the full, original model (Matlab implementation). The dataset used in the comparison is a set of 50 MS2 fluorescence traces of the ush gene in a *Drosophila* embryo, where active transcription occurs during a 30 minute time window. A window size of 19 is required to model the fluorescence traces. The curve plotted in blue shows the result of increasing the window size upon the computational time required for a single expectation-maximisation step for the full model; the exponential scaling of the algorithm with window size is apparent. The computational time for the truncated model (red, *M* = 128 compound states, 90s per step) is essentially de-coupled from window size / gene length, allowing for application of the truncated model to a much wider set of window sizes (Figure 4(C)). For short genes, the original version model is faster due to less computational overhead associated with truncation, e.g. calculating and eliminating least likely states etc. The benefits of the truncated version of the model become apparent at longer gene lengths, where exponentially increasing computation time makes inference impractical. A window size of 30+ may be needed for both much longer *Drosophila* genes and vertebrate genes, making use of the full model infeasible.

### 3.3 Application of the algorithm to real data

An example of using the model to infer single-cell parameters is shown in Figure 5, using example data from Hoppe *et al.* (2020) (different embryo to that highlighted in the original paper). The aim of the paper was to use the parameters inferred by *burstInfer* to investigate regulatory control of Bone Morphogenetic Protein (BMP) target genes in the early Drosophila embryo, focussing on dorsal-ventral patterning of the early embryo. MS2 imaging was used to generate movies of transcriptional activity of one of the BMP target genes studied in the paper, *ush*, during nuclear cycle 14. The expression domain of *ush* forms a broad stripe down the anterior-posterior axis on the dorsal side of the embryo (Ashe *et al.*, 2000), which mirrors the expression levels of the BMP Decapentaplegic (Dpp) (Figure 5A) (Bier and De Robertis, 2015; Deignan *et al.*, 2016; Eldar *et al.*, 2002; Umulis *et al.*, 2010). Cells falling within the Dpp gradient express Dpp target genes in a concentration-dependent manner - intermediate levels of signalling are sufficient to activate ush, for example.

To investigate spatial regulation of Dpp target genes, MS2 movies were recorded in the embryo during nuclear cycle 14. Each embryo was divided into three separate regions corresponding to different signalling levels, determined by either distance from the midline or through the use of a clustering-based approach. *burstInfer* was then trained on each of these three regions, giving estimates of *k_on_, k_off_* and Pol II loading rate (emission) for each section of the embryo. These regional parameters were then used to infer single-cell parameters (Figure 5B) and promoter traces (Figure 5C) for each cell within the expression domain. Figure 5B shows heatmaps of mean expression and three example single-cell parameters for *ush* - the region shown here represents a subset of the expression domain shown in the cartoon in Figure 5A. Mean expression corresponds to the mean recorded fluorescence for each cell, with the arbitrary fluorescence signals converted into number of Pol II. The single cell occupancy, *k_on_ and k_off_* parameters were calculated using *burstInfer*. From these heatmaps the strong similarity between mean expression and occupancy is immediately apparent, along with the slightly weaker similarity between expression levels and *k_on_* (Figure 5B). In order to quantify the dependency of expression levels on each of these three parameters (along with other derived parameters, such as burst duration and frequency), correlation analysis was carried out on the single-cell expression data and inferred parameters.

This analysis revealed a very strong correlation between expression levels and occupancy, with effectively no correlation between expression and *k_off_* (Hoppe *et al.*, 2020). Pol II loading rate (the HMM emission parameter) was flat across the expression domain Figure (5B). As occupancy depends upon both *k_on_* (which did exhibit strong correlation) and *k_off_*, the results indicated that expression levels were regulated through modulation of *k_on_*, the promoter activation rate. Representative single cell fluorescence and promoter traces for each region show that nuclei experiencing high signalling produce more transcriptional bursts compared to other regions (Figure 5C). The single-cell parameters extracted from quantification of traces like these were used to create the heatmaps shown in Figure 5B. Code to re-create these figures is included in the *burstInfer* GitHub repository.

## 4 Discussion

We have presented an algorithm for efficient inference of transcriptional kinetic parameters, with the aim of improving upon an existing memory-adjusted Hidden Markov model (Lammers *et al.*, 2020) by reducing the computational time required for inference. We also provide methods for inferring the parameters of transcriptional dynamics in single cells. The algorithm allows for the inference of burst amplitude, duration and frequency from MS2 data, which we expect to be of interest to researchers working on transcriptional regulation. The MS2-MCP system has provided researchers with high-quality data relating to transcriptional activity in individual cells, and has been used to provide insight into the dynamics of transcription. However, the persistence present within the MS2 signal presents a challenge when attempting to infer kinetic parameters using these particular datasets. Our algorithm allows efficient inference of kinetic parameters for longer genes than is currently possible.

A comparison of the running time for a single step of the expectation maximisation algorithm for both the full and truncated models demonstrated the reduction in computational time while using the truncated model on the *Drosophila* gene *ush*, which would require a window size of 19 for inference. The time taken for a single expectation-maximisation step at window size 19 (42 minutes) would render inference using the full model for this particular gene computationally infeasible, particularly if repeated likelihood computations, e.g. for statistical approaches such as bootstrapping or MCMC sampling, are required. The truncated model, in comparison, does not scale significantly with gene length and is instead primarily limited by a linear dependence on the size of the training dataset. This ability to model genes of arbitrary length should allow the model to be applied to more complex organisms, with longer genes, than *Drosophila*.

A demonstration of applying the model to infer transcriptional parameters in *Drosophila* was outlined in (Hoppe *et al.*, 2020). In that study, *burstInfer* was used to investigate regulation of BMP target genes in the early embryo through dividing embryos into regions corresponding to different BMP signalling levels then training the model on the MS2 datasets for each of these regions. The fitted model was used to generate transcriptional parameters in single cells, allowing the investigation of spatial changes in bursting dynamics. We expect that the method would be well suited for analysis of similar datasets in other systems.

## Acknowledgements

The authors would like to thank Catherine Sutcliffe for her help with the MS2-MCP experiments used to generate the datasets shown in this paper.

## Funding

This work has been su
pported by the Wellcome Trust [Grant Refs. 204832/B/16/Z, 204832/Z/16/Z, 215187/Z/19/Z].

## Notes

https://github.com/ManchesterBioinference/burstInfer

### Competing Interest Statement

The authors have declared no competing interest.

## References

Ashe, H.L. et al. (2000) Dpp signaling thresholds in the dorsal ectoderm of the Drosophila embryo, Development, 127, 3305–3312

Bahar Halpern, K. et al. (2015) Bursty gene expression in the intact mammalian liver, Mol. Cell, 58, 147–156

Berrocal, A. et al. (2018) Kinetic sculpting of the seven stripes of the Drosophila even-skipped gene, bioRxiv, 1, 335901*

Bertrand, E. et al. (1998) Localization of ASH1 mRNA Particles in Living Yeast, Mol. Cell, 2, 437–445

Bier, E. and De Robertis, E.M. (2015) BMP gradients: A paradigm for morphogen-mediated developmental patterning, Science, 348, aaa5838

Bothma, J.P. et al. (2014) Dynamic regulation of eve stripe 2 expression reveals transcriptional bursts in living Drosophila embryos, Proc. Natl. Acad. Sci., 29, 10598–10603

Chubb, J.R. et al. (2006) Transcriptional Pulsing of a Developmental Gene, Current Biology, 16, 1018–1025

Corrigan, A.M. et al. (2016) A continuum model of transcriptional bursting, Elife, 5, 1–38

Coulon, A. et al. (2013) Eukaryotic transcriptional dynamics: From single molecules to cell populations, Nat. Rev. Genet., 14, 572–584

Dar, R.D. et al. (2012) Transcriptional burst frequency and burst size are equally modulated across the human genome, PNAS, 109, 17454–17459

Deignan, L. et al. (2016) Regulation of the BMP Signaling-Responsive Transcriptional Network in the Drosophila Embryo, PLoS Genetics, 12, e1006164

Eldar, A. et al. (2002) Robustness of the BMP morphogen gradient in Drosophila embryonic patterning, Nature, 419, 304–308

Eldar, A. et al. (2010) Functional roles for noise in genetic circuits, Nature Reviews, 467, 167–173

Fukaya, T., et al. (2016) Enhancer Control of Transcriptional Bursting, Cell, 166, 358–368

Garcia, H.G. et al. (2013) Quantitative imaging of transcription in living Drosophila embryos links polymerase activity to patterning, Curr. Biol., 23, 2140–2145*

Golding, I. et al. (2005) Real-time kinetics of gene activity in individual bacteria, Cell, 123, 1025–1036

Gómez-Schiavon, M. et al. (2017) BayFish: Bayesian inference of transcription dynamics from population snapshots of single-molecule RNA FISH in single cells, Genome Biol., 18, 164*

Gregor, T. et al. (2014) The embryo as a laboratory: Quantifying transcription in Drosophila, Trends Genet., 30, 364–375

Hoppe, c. et al. (2020) Modulation of the Promoter Activation Rate Dictates the Transcriptional Response to Graded BMP Signaling Levels in the Drosophila Embryo, Dev. Cell, 54, 727–741

Lammers, N. et al. (2020) Multimodal transcriptional control of pattern formation in embryonic development PNAS, 117(2), 836–847

Lenstra, T.L. et al. (2016) Transcription Dynamics in Living Cells, Annu. Rev. Biophys., 45, 25–47

Li, C. et al. (2018) Frequency Modulation of Transcriptional Bursting Enables Sensitive and Rapid Gene Regulation, Cell Syst., 6, 409–423

Lucas, T. et al. (2013) Live Imaging of Bicoid-Dependent Transcription in Drosophila Embryos, Curr. Biol., 23, 2135–2139

Mueller, F. et al. (2013) Automatic counting of transcripts in 3D FISH images, Nat. Methods, 10, 277–278

Pichon, X. et al. (2018) A Growing Toolbox to Image Gene Expression in Single Cells: Sensitive Approaches for Demanding Challenges, Mol. Cell, 70, 468–480

Raj, A. et al. (2006) Stochastic mRNA synthesis in mammalian cells, PLoS Biol., 4, 1707–1719

Raj, A. et al. (2008) Nature, Nurture, or Chance: Stochastic Gene Expression and Its Consequences, Cell, 135, 216–226*

So, L.H. et al. (2011) General properties of transcriptional time series in Escherichia coli, Nat. Genet., 43, 554–560

Tantale, K. et al. (2016) A single-molecule view of transcription reveals convoys of RNA polymerases and multi-scale bursting, Nat. Commun., 7, 12248*

Umulis, D.M. et al. (2010) Organism-Scale Modeling of Early Drosophila Patterning via Bone Morphogenetic Proteins, Dev. Cell, 18, 260–274

Zoller, B. et al. (2018) Diverse Spatial Expression Patterns Emerge from Unified Kinetics of Transcriptional Bursting, Cell, 175, 835–847*

